# Illuminating the dark depths inside coral

**DOI:** 10.1101/534842

**Authors:** Chichi Liu, Shuk Han Cheng, Senjie Lin

**Affiliations:** State Key Laboratory of Marine Environmental Science, College of Ocean and Earth Sciences, Xiamen University, Xiamen, China; Department of Biomedical Sciences, City University of Hong Kong, Hong Kong, China; Department of Marine Sciences, University of Connecticut, Groton, CT, USA

**Keywords:** Coral, Symbiodiniaceae, Tissue clearing, Light sheet fluorescence microscopy, 3D imaging, Cellular hypoxia

## Abstract

The ability to observe in situ 3D distribution and dynamics of endosymbionts in corals is crucial for gaining a mechanistic understanding of coral bleaching and reef degradation. Here, we report the development of a tissue clearing (TC)-coupled light sheet fluorescence microscopy (LSFM) method for 3D imaging of the coral holobiont at single-cell resolution. The initial applications have demonstrated the ability of this technique to provide high space-resolution quantitative information of endosymbiont abundance and distribution within corals. With specific fluorescent probes or assays, TC-LSFM also revealed spatial distribution and dynamics of physiological conditions (such as cell proliferation, apoptosis, and hypoxia response) in both corals and their endosymbionts. This tool is highly promising for in situ and in-depth data acquisition to illuminate coral symbiosis and health conditions in the changing marine environment, providing fundamental information for coral reef conservation and restoration.

## Introduction

Coral reefs, the most species-rich and productive ecosystem in the ocean, are threatened by accelerated global climate and environmental changes [1,2,3,4,5,6]. Damages are mostly rooted in the disruption of the symbiosis between hermatypic corals and their endosymbionts, which are algae belonging to the dinoflagellate family Symbiodiniaceae [7,8,9]. Coral beaching, a major cause of coral degradation, occurs as the coral host expels the endosymbionts and hence loses the characteristic pigment of the algae in response to environmental stress [10,11]. To understand how environmental stress causes coral bleaching and why different species of Symbiodiniaceae respond differently to stress, researchers performed extensive studies from molecular, cellular, to ecosystem levels [11,12,13,14,15,16]. Live observation of corals and documentation of their behavior or physiology have also been carried out through underwater microscopy [17] and coral-on-a-chip approach [18]. However, spatial organization of the endosymbionts and spatially resolved physiological dynamics of the holobiont in the process of stress response are largely intractable due to the lack of proper techniques that allow in-depth observations in toto.

Here, leveraging the power of light sheet fluorescent microscopy (LSFM) and tissue clearing technology (TC), we developed a whole-mount 3D imaging method to directly observe symbiont organization and coral/symbiont physiologies in toto. LSFM features a fast acquisition speed, a low phototoxicity, and a high efficiency for imaging thick biological samples [19,20,21,22]. It alone is promising for imaging fixed [23] and live [24] coral samples, but the resolution and imaging depth are compromised by the blockage of illumination by coral skeleton and the low transparency of coral tissues. Tissue clearing, a frequently employed tool in biomedical research [25,26,27,28], has the potential to address this issue. In this study, we developed a protocol combining LSFM and TC. In the initial limited-scope application of this method, we successfully obtained 3D see-through images of coral tissue and demonstrated a broad range of utility in studying physiologies and stress responses of coral hosts and their endosymbionts.

## Materials and methods

### Coral sample collection and maintenance

Healthy *Seriatopora hystrix* and *Seriatopora* sp. were acquired from a coral aquarium tank. Bleached *S. hystrix* and *S. stellate* were collected by SCUBA diving in Bohol, Philippines. Similarly, bleached *Acropora* sp. were collected by SCUBA diving in Hong Kong. Coral samples were fixed in 4% paraformaldehyde prepared in 0.45 μm filtered seawater (FSW) immediately after collection and processed in the laboratory.

### Tissue clearing of coral sample

The fixed coral fragments were washed with 0.01 M PBS thrice at room temperature with gentle shaking. Coral skeleton was removed through decalcification conducted in 10% EDTA at pH 7.4. The solution was replaced twice every day until the whole skeleton was removed. Methanol was used for decolorizing and permeabilization. The samples were washed with PBS thrice and dehydrated in 50%, 75%, 90%, and 100% methanol in 0.01 M PBS for 10 min at each step with gentle shaking. The samples were then rehydrated in 90%, 75%, and 50% methanol in PBST (0.01 M PBS containing 0.2% Triton X-100) for 10 min at each step with gentle shaking. Incubation time was carefully controlled to preserve the auto-fluorescence signal of chlorophyll during LSFM imaging. The samples were washed with PBST thrice and fluorescence labeled in accordance with the detailed procedure as described below. The samples for measuring the density of Symbiodiniaceae only were directly subjected to refractive index (RI) matching step. RI matching was performed by immersing the sample in a RI matching solution (RIMS), which was obtained by dissolving 20 g of Nycodenz (Axis-Shield Diagnostics) in 16 mL of 0.02 M PBS and adjusting RI to 1.45 by using water. After 2 h to 4 h of immersion, RIMS was renewed to maintain the RI value. DAPI was added to RIMS in the second round of immersion with the final concentration of 1 μg/mL to improve the staining of the coral tissue. After 1 h to 2 h of immersion and staining, the cleared coral sample was ready for LSFM imaging as described below.

### Micropropagation of *P. damicornis*

Coral micropropagates were obtained through a polyp bailout assay. *P. damicornis* fragments were placed in a Petri dish filled with FSW, the salinity of which was gradually adjusted to 50 PSU within 5 h by using high-salinity FSW (60 PSU). The bailout polyps were examined under a stereomicroscope. Some of the micropropagates were fixed immediately with 4% paraformaldehyde prepared in FSW. These samples were used as newly separated polyps for apoptotic cell detection via a TUNEL assay. Some of the micropropagates were gradually transferred to normal salinity at 33–35 PSU and frequently examined under a stereomicroscope. Once the micropropagates flattened, which indicated the initiation of reattachment, the samples were fixed as previously described. These samples were used as newly reattached coral micropropagates for cell mitosis detection through immunofluorescence assay.

### Cell proliferation and apoptosis detection

Fluorescence labeling of proliferating cells were achieved by using an EdU cell proliferation assay kit (C10638, Thermo Fisher Scientific, US). A coral fragment was placed in a sealed chamber filled with 50–100 times volume of FSW containing 20 μM EdU. The chamber was incubated in a coral aquarium tank to maintain an optimal temperature of 26 °C and a light intensity of approximately 100 μE m^−2^ s^−1^. After 1 h of incubation, the coral branch was cut into small pieces by using a bone cutter, fixed in 4% paraformaldehyde prepared in FSW, decalcified, and decolorized as previously described. EdU was detected in accordance with the manufacturer’s instructions with slight modifications. In general, the volumes of the reagents used in the assay were adjusted to about 20–30 times the volume of the coral samples, and they were incubated two times longer to achieve the whole tissue detection of the incorporated EdU. The samples were then subjected to DAPI staining and RI matching as previously described. Light exposure was avoided.

TUNEL assay (G3250, Promega, US) was used for fluorescence labeling of apoptotic cells. Sample fixation, decalcification, and decolorization were conducted as described above. The samples were permeabilized with 20–30 times the volume of 20 μg mL^−1^ Proteinase K at room temperature for 30 min, washed with 0.01 M PBS thrice, and equilibrated at room temperature for 20 min with 20 times the volume of TdT buffer. A TdT reaction was carried out with 20 times the volume of a TdT reaction mix at 37 °C for 90 min. TdT enzyme negative control was conducted simultaneously. After the TdT reaction, the samples were subjected to DAPI staining and RI matching as described above.

### Whole mount immunofluorescence of histone H3 phosphorylation in coral tissues

For immunofluorescence staining, sample fixation, decalcification, and decolorization were conducted as described earlier. Blocking was performed using blocking buffer (PBST containing 2% BSA and 2% goat serum) at room temperature for 2 h to 4 h. Primary antibody (rabbit anti-histone H3 [Ser10]; EMD Millipore, USA) was diluted (1:500) in blocking buffer and incubated with the coral tissues at 4 °C overnight. The coral sample was then washed with PBST thrice for 30 min at each step and incubated with Alexa Fluor 488-conjugated goat anti-rabbit IgG (A11034, Invitrogen, 1:200, Invitrogen) as a secondary antibody at 4 °C overnight. The samples were washed thrice with PBST for 30 min at each step and subjected to DAPI staining and RI matching as described above.

### Detection of cellular hypoxia of coral tissues

A coral nubbin was placed in a sealed respiration chamber filled with FSW in the dark to induce coral tissue hypoxia. After 5 h of incubation, pimonidazole hydrochloride (Hypoxyprobe-Red549 Kits, Hypoxyprobe, USA) was added with a final concentration of 300 nM. After 1 h of incubation, coral nubbins were then fixed with 4% PFA in FSW for 2 h at room temperature or overnight at 4°C. The whole process was conducted under low light to avoid changes in cellular oxygen concentrations due to photosynthesis. The samples were then decalcified and decolorized as described earlier. Dylight™549 conjugated antibody (1:50 dilution) was utilized to detect the intracellular content of a pimonidazole adduct. The subsequent procedure was similar to the whole mount immunofluorescence protocol described above. The coral samples that were not subjected to hypoxyprobe incubation were used as negative control, while the coral samples incubated in the dark with air bubbles were set as normoxic control. The samples were finally subjected to DAPI staining and RI matching as described earlier.

### Coral sample mounting and light sheet imaging

3D images of the coral samples were taken using ZEISS lightsheet Z.1 system. The samples were mounted in accordance with the guidelines provided by ZEISS with some modifications. A glass capillary with an inner diameter of 2 mm, a 1 mL syringe with an inner diameter of 4.7 mm, and a glass tube with inner diameters of 3.5 and 6 mm with a custom-made plunger were used to mount the coral samples with an appropriate size. A 3D-printed adaptor was used to fasten the glass tube to the syringe sample holder of the microscope. After RI was adjusted, the coral samples were transferred to a mounting medium (RIMS with 1.5% low-melt agarose, RI = 1.450, 50 °C to 55 °C), gently inverted several times to avoid bubble formation, and incubated for 5 min. The sample was then loaded to the mounting tube with an appropriate inner diameter and placed at 4 °C in a light-proof moisture chamber for at least 1 h to allow the solidification of the mounting medium. The samples were mounted and immediately imaged or stored at 4 °C in the moisture chamber until they were subsequently imaged. Before imaging was performed, the samples stored for hours or longer were re-immersed in RIMS to re-adjust RI to 1.45.

A clearing (RI = 1.45) optical setting with a 5× objective was applied to the ZEISS lightsheet Z.1 system, whereas a 0.6–1.5 × tube lens was used depending on the size of the coral sample. Under this condition, z-stepping was 2 or 2.5 μm with 4 or 5 μm thickness of a laser sheet. In this setting, 500 or 400 optical sections for 1 mm-thick coral tissues were produced, thereby allowing the precise 3D reconstruction of the coral sample. The imaging chamber was filled with RIMS (RI = 1.45) and frequently monitored using a refractometer (NAR-1T, ATAGO, US). The cylinder of the solidified mounting medium containing the coral sample was pushed out from the glass capillary or the syringe, equilibrated for 10 min to 20 min, and imaged. The coral samples with a thickness of >600 μm were subjected to two-view image acquisition and post-imaging fusion. DAPI was excited using a 410 nm laser and the fluorescence signal was collected with a 490 nm bandpass filter. Coral green fluorescence protein and apoptosis TUNEL assays were excited using a 488 nm laser and collected with a 533 nm bandpass filter. Dylight™549, Alexa Fluor 555, and CY3 were excited using a 561 nm laser and collected with a 590 nm bandpass filter. The autofluorescence of the chlorophyll was excited using a 410 nm laser and collected with a 610 nm long-pass emission filter. The samples in one experiment were captured under identical conditions of laser power, exposure time, laser sheet thickness, and z-stepping.

### Quantitative analysis of 3D image data

Multi-view image data were fused and deconvoluted with ZEN software (Black edition, ZEISS). Identical gamma value and dynamic range were applied to all of the samples to obtain consistent results. Afterward, 3D reconstruction was performed using arivis Vision4D (version 2.12.4). The volume of the chlorophyll autofluorescence signal of Symbiodiniaceae cells was calculated using an analysis pipeline of the intensity threshold and followed by a segment filter in Vision4D. All of the signals were sorted in the order of volume first, and the roundness of x-y projection and the sphericity of the sorted fluorescence signal were plotted in the same order. The drop of roundness and sphericity indicated that the signal contained two or more Symbiodiniaceae cells that were too close to be separated (Supplementary Figs. 2e and f). Based on the signal volume value and information on roundness and sphericity, single-cell signals were identified. The average volume of the signal from a single Symbiodiniaceae cell was then calculated from the single-cell signals. The total number of Symbiodiniaceae cells was then obtained by dividing the total signal volume with the average single-cell signal. The average diameter of Symbiodiniaceae cells was calculated from the area of the x-y projection of the single Symbiodiniaceae signal. The number of signals of the other physiological parameters, such as cell proliferation and cell apoptosis, were counted directly by using a similar analysis pipeline with a proper threshold setting.

**Fig. 1.**
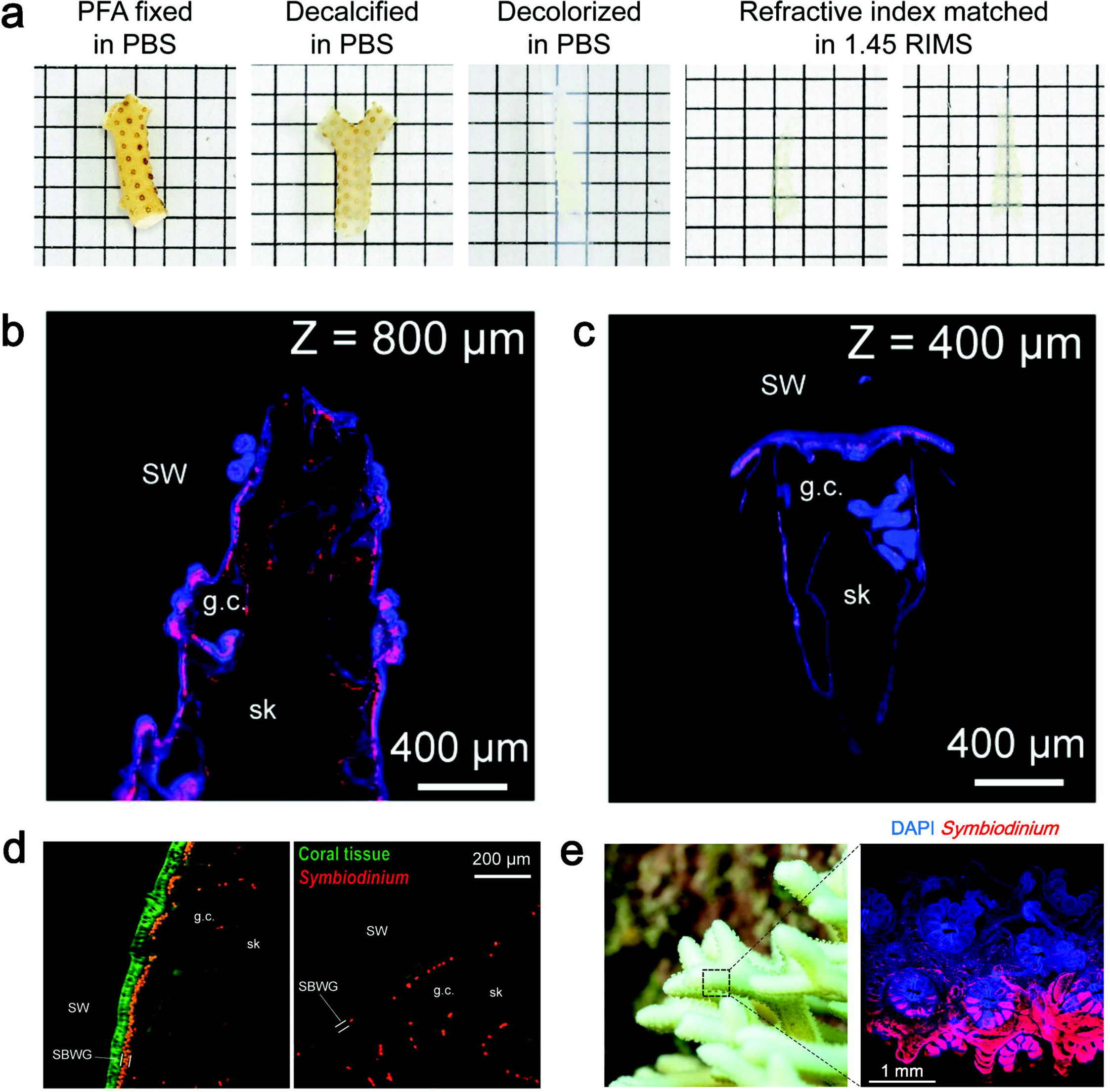
Clearing and light sheet imaging of the coral sample. **a** Brightfield images demonstrating the transparency of the coral samples after each step of the treatment. **b** Image illustrating 1 out of 857 optical sections of branch tip imaging (z = 800 μm; for more data, see Supplementary Fig. 1a and Supplementary movie 1). **c** Image showing 1 out of 635 optical sections of the isolated polyp of *Seriatopora* sp. (z = 400 μm; for more data, see Supplementary Fig. 1b). **d** Optical section of unbleached (left panel) and bleached (right panel) *Acropora* sp․. Green indicates the fluorescence of the green fluorescence protein in the coral tissue. **e** Close-up image of coral *S. stellate* with light-induced bleaching on the upper surface (left panel) and maximum intensity projection after FLSM-TC imaging (right panel). SBWG, surface body wall gastrodermis; SW, seawater; sk, decalcified skeleton; g.c., gastric cavity.

**Fig. 2:**
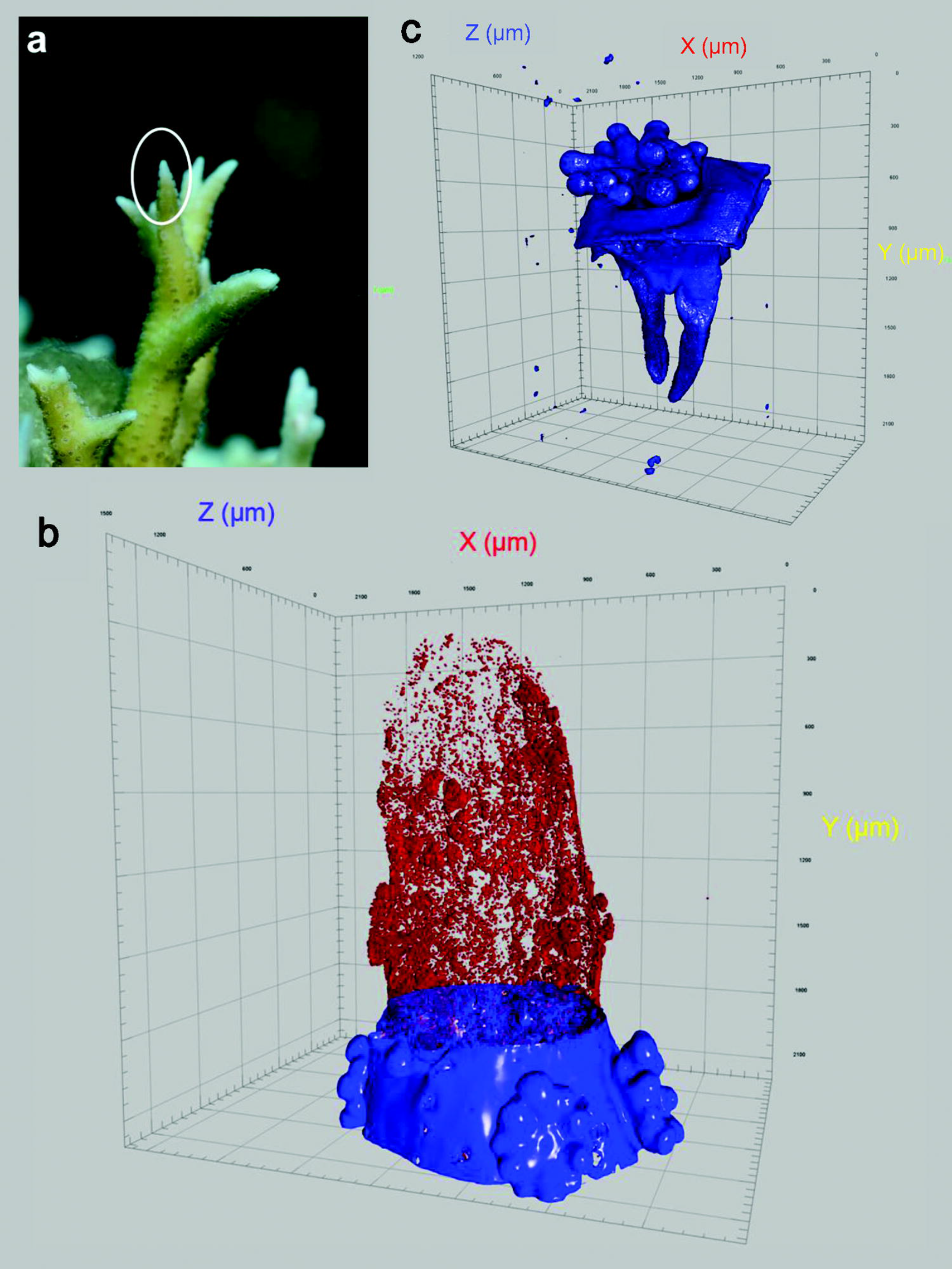
3D reconstruction of the coral sample based on serial optical sections. **a** Close-up image of coral *S. hystrix* with a branch tip (circled) to be imaged. **b** 3D reconstruction of the *S. hystrix* branch tip showing the distribution pattern of Symbiodiniaceae recognizable from its red autofluorescence of chlorophylls (shown in the upper part of the image and obtained when the coral tissue was digitally removed to create a see-through effect; blue fluorescence of the lower part is from DNA-DAPI stain of the un-removed coral tissue). For more data, see Supplementary Movie 2. **c** 3D reconstruction of a *Seriatopora* sp. polyp (for more data, see Supplementary Movie 3).

DAPI signal was used to calculate the volume and surface area of the coral samples. Serial longitudinal optical sections were converted to cross-sections by using the series modify function of ZEN software and then fed to Vision4D for 3D reconstruction to calculate volume. The intensity threshold and the segment filter were used as an analysis pipeline, and the same threshold settings were applied to all of the samples in one experiment. The surface areas of the branch coral samples were estimated as the area of the equivalent unfolded cylinder based on sample height and volume. The scope of analysis was adjusted in three dimensions to calculate the number of Symbiodiniaceae cells and the volume of the tissues in a specific area.

## Results

### See-through 3D imaging of the coral samples

The coral samples were fixed on site by using paraformaldehyde to preserve under in situ conditions. Decolorization was conducted to the samples while the fluorescent signal of the chlorophyll was still detectable to easily distinguish Symbiodiniaceae from the coral tissue and to minimize light absorption. The transparency of the coral sample immersed in an aqueous tissue-clearing solution with a RI of 1.45 was significantly enhanced (Fig. 1a). Target-specific information could be obtained through bioimage informatics analysis by collecting the *in toto* fluorescence signal of the tissue-cleared coral sample through LSFM. As an example of our initial applications of the technique, the serial optical sections of a branch tip of *S. hystrix* were collected, and each section showed a clear fluorescence image at a certain depth of the sample (Fig. 1b, Supplementary Fig. 1a, and Supplementary Movie 1). The single polyps of *Seriatopora* sp. were successfully separated from the decalcified tissue and imaged, and optical sections deep in the tissues were acquired to clearly show the structure of the gastric cavity and the organs inside, such as the mesenterial filament (Fig. 1c and Supplementary Fig. 1b). With these methods, the coral tissues could be clearly visualized through the blue fluorescence of the DNA-DAPI stain and the populated and heterogeneously distributed Symbiodiniaceae through the overwhelming red autofluorescence of chlorophylls.

The TC-LSFM imaging of the healthy and bleached *Acropora* sp. samples revealed that the Symbiodiniaceae density decreased in the surface body wall gastrodermis (SBWG) but not in the aboral tissue (Fig. 1d). With the high spatial resolution of the technique, localized bleaching, such as that induced by excess light on the upper side of *S. stellate* branch, could be examined and different Symbiodiniaceae densities between tentacles of one coral polyp could be documented (Fig. 1e). The 3D reconstruction from the numerous optical sections yielded the high accuracy and fine spatial resolution of Symbiodiniaceae distribution in the 3D space within the coral branch tips and within the single polyps (Fig. 2; Supplementary Movies 2 and 3).

### Quantitative analysis of Symbiodiniaceae 3D distribution

The fluorescent signals of Symbiodiniaceae were easily identified in the 3D-rendered volumetric image of a coral sample. However, the direct counting of discrete signals would seriously underestimate the number of Symbiodiniaceae cells because multiple Symbiodiniaceae cells might be counted as one when they were too close to one another to be separated (e.g., polyp tentacles). An image informatics workflow was developed to address the challenge based on the concept of the signal volume (Fig. 3a). In some areas of the coral sample, the boundary of the cell signals could be easily identified because the cells existed singly or were well separated. In most parts of the coral sample, the boundary was unidentifiable (Fig. 3b). In such a sample, the 2D projections of the signals from single cells (small signal volumes) were perfectly round, whereas the 2D projections of the signals from two or more clumped Symbiodiniaceae cells (signal volume exceeding a certain value) showed a rapidly decreased roundness (Fig. 3c). The identified clump of the multiple cells could be confirmed by the same trend of the sphericity of the 3D-rendered fluorescence signals (Fig. 3d). Thus, with the help of the roundness and sphericity information, the signals of the single cells could be accurately identified. The Symbiodiniaceae cell count in a cell clump could be obtained by dividing the total Symbiodiniaceae fluorescence signal volume with the single cell signal volume on the basis of the average signal volume of a single Symbiodiniaceae cell. The volume or surface area of the coral sample required for cell density estimation could be calculated from rendered volumetric image. Although DAPI stains DNA only in the nuclei, the fluorescence signal tended to cover the entire tissue surface of the coral sample because of the high cell density in the coral epidermis and could hence be used to reconstruct the coral tissue surface area. In the branch tip shown as an example (Fig. 1b), our calculation revealed that the Symbiodiniaceae density of *S. hystrix* ranged from 6.6×10^6^ cells cm^−2^ at the tip to 13.5×10^6^ cells cm^−2^ toward the base (Fig. 3e). The conventional method would only generate an average Symbiodiniaceae density (~10^7^ cells cm^−2^ assuming 100% cell recovery) for the whole branch segment.

**Fig. 3:**
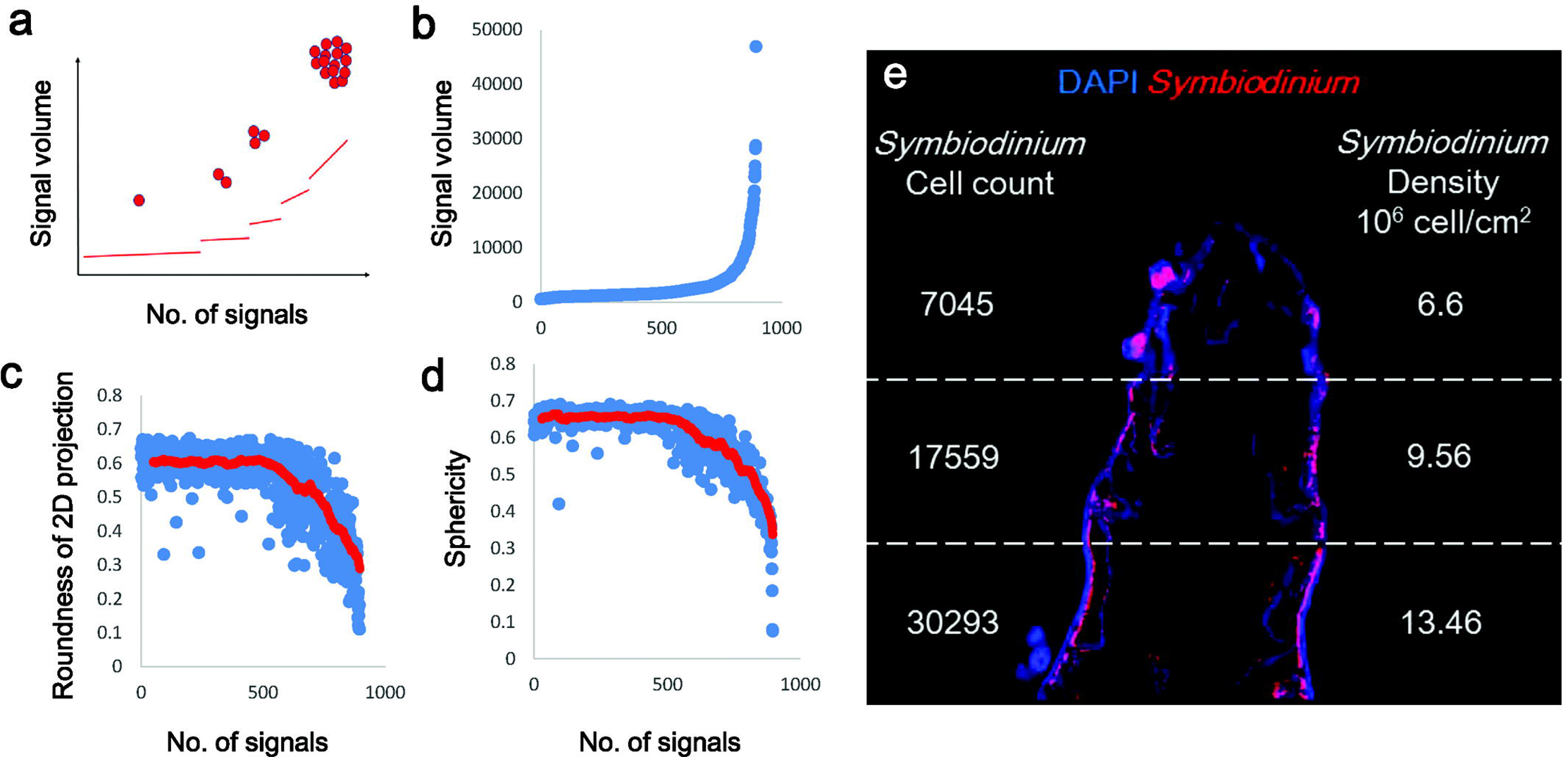
Quantification of Symbiodiniaceae density based on 3D image data by using the signal volume algorithm. **a** Concept of the signal volume calculation of a single Symbiodiniaceae cell. Filled circles represent a single cell or groups of cells. Lines depict signal volumes corresponding to the number of signals represented by the number of cells (filled circles). **b** Volumes of the identified Symbiodiniaceae signals in a coral sample increased sharply as Symbiodiniaceae cells clumped together. **c** Roundness of the 2D projection of the Symbiodiniaceae signal, which dropped precipitously when some Symbiodiniaceae cells clumped together and the signal volume exceeded a certain level. **d** Sphericity of the rendered 3D Symbiodiniaceae fluorescence signal, which declined precipitously when some Symbiodiniaceae cells clumped together and the signal volume exceeded a certain level. **e** Quantification of the number and density (cells/cm^2^) of Symbiodiniaceae cells in the *S. hystrix* branch tip in Fig. 1b as estimated using the signal volume algorithm by dividing the total signal volume with the single-cell signal volume. Single-cell signal was identified with the aid of the roundness and sphericity information. In all of the images, blue indicates the fluorescence of DNA-DAPI stain, whereas red denotes the autofluorescence of chlorophylls in Symbiodiniaceae cells.

### 3D imaging of the coral physiology

TC-LSFM could also be performed to visualize the physiological conditions in the coral tissues. Using EdU cell proliferation assay, we imaged the numbers and spatial patterns of proliferating cells in a coral branch tip, showing a high density of proliferating cells and a low density of Symbiodiniaceae cells in the apex (Fig. 4a). We obtained *Pocillopora damicornis* micropropagates through a “bailout” procedure and conducted a TUNEL assay to detect apoptotic cells during bailout and reattachment. Numerous apoptotic cells were observed, suggesting that tissue remodeling occurred immediately after the polyp bailed out (Fig. 4b). We also performed the whole mount fluorescence immunolabeling of histone phosphorylation, which is a specific marker of mitosis, in newly reattached coral micropropagates. Mitosis was quickly initiated after re-attachment in the coral (Fig. 4c) but not in the unattached polyp (data not shown). Although the coral and the Symbiodiniaceae cells were labeled at the same time, the signal was distinguishable by using a segment filter because of their different volumes and red fluorescence signals from chlorophylls in Symbiodiniaceae. The mitotic status of the labeled Symbiodiniaceae cells was also evidenced by their increased cell size (data not shown). The 3D reconstruction also showed that the polyp flattened after re-attachment and the distribution of Symbiodiniaceae (Supplementary Movie 4). To extend the application of this method to coral reproduction research, we also collected the 3D images of spermary of *Acropora* sp., and the developing lumen indicated that the spermary was at stage III of development (Fig. 4d and Supplementary Movie 5).

**Fig. 4:**
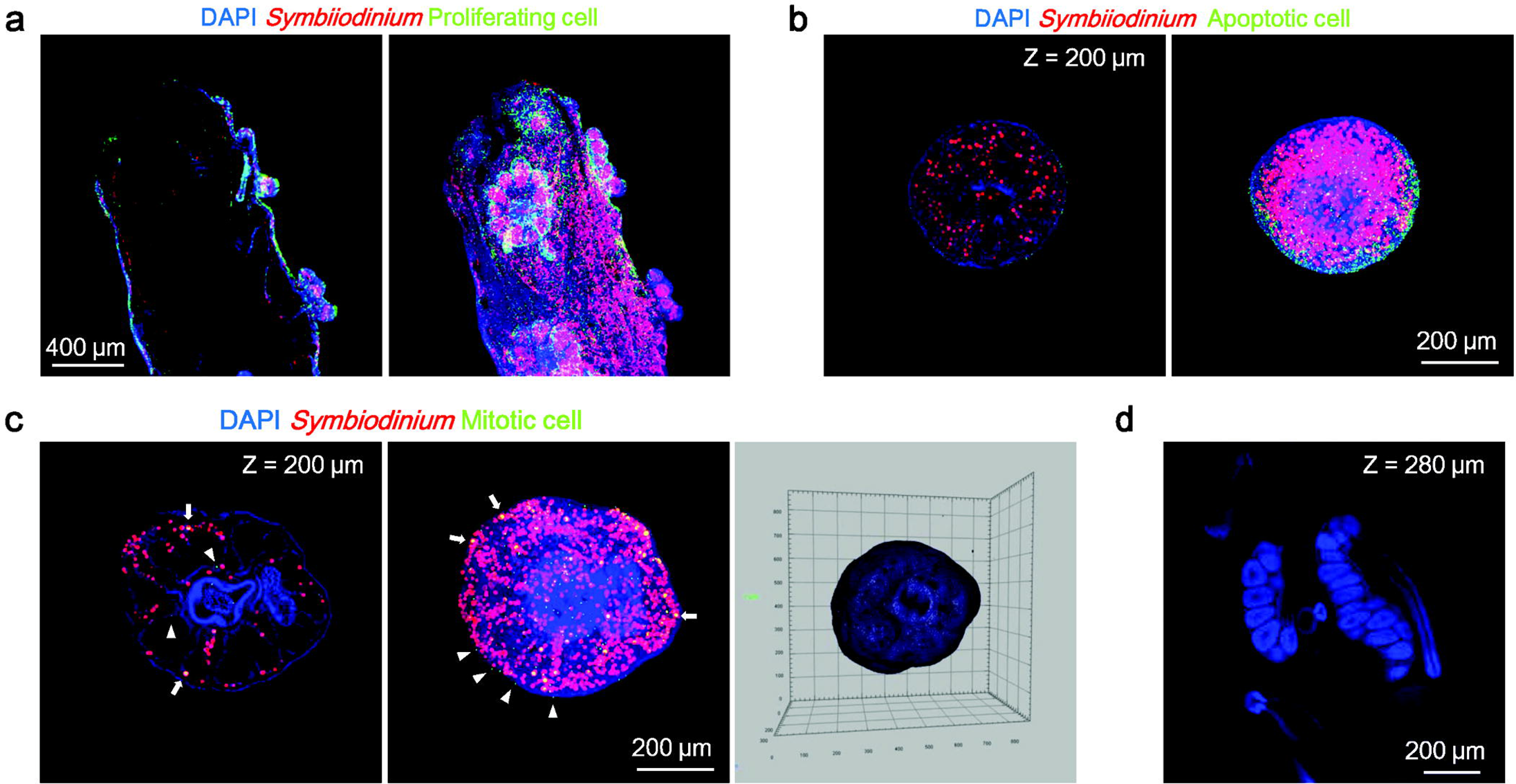
TC-LSFM coupled with various fluorescence labeling assays to probe the physiologies of coral-Symbiodiniaceae systems. **a** Optical section at the *z* depth of 600 μm (left panel) and the maximum intensity projection (right panel) of *Seriatopora* sp. branch tip showing the proliferating cells detected by using EdU cell proliferation assay (green fluorescence). **b** Optical section at the *z* depth of 200 μm (left panel) and the maximum intensity projection (right panel) of apoptotic cells detected by using TUNEL assay (green fluorescence) in the bailout polyp of *Pocillopora damicornis*. **c** Immunolabeling of mitotic coral cells (arrowheads) and Symbiodiniaceae cells (arrows) detected using the histone H3 phosphorylation-specific antibody in the reattached bailout polyp of *P. damicornis*, left: single optical section (z = 200 μm); middle: maximum intensity projection; right: 3D reconstruction of the reattached polyp (for more data, see Supplementary Movie 4). **d** Optical section of *Acropora* sp. at the *z* depth of 280 μm showing the clear structure of spermary (3D reconstruction of the sample shown in Supplementary Movie 5). In all images, blue indicates the fluorescence of DNA-DAPI stain, whereas red denotes the autofluorescence of chlorophylls in Symbiodiniaceae cells.

### Detection of cellular hypoxia in coral holobionts

Coral cells can experience hypoxia and severe stress at night due to the on and off dynamics of photosynthetic oxygen production [29]. To address the challenge in detecting cellular hypoxia in a coral holobiont, we applied a hypoxyprobe, which is widely used in biomedical and clinical science. A pimonidazole-based hypoxyprobe easily diffuses into cells and forms adducts when the intracellular oxygen level decreases below 10 mmHg and allows post-fixation detection through immunofluorescence. Our results showed that the cellular hypoxia of the coral and the Symbiodiniaceae cells could be detected using TC-LSFM method. In this experiment, coral cell hypoxia occurred in the aboral tissue but not in the epidermal layer under the dark condition without an oxygen supply. A gradient of hypoxyprobe signals (green fluorescence) was observed along the tissue depth (Fig. 5a). No fluorescence signal was detected in the pimonidazole negative control, whereas only a weak signal was detected in the tentacle area of the normoxic control (dark condition with oxygen supply) (Fig. 5a). Some of the Symbiodiniaceae cells that were spatially unevenly distributed showed signs of cellular hypoxia (Fig. 5b). The percentage of the hypoxia-stressed Symbiodiniaceae cells could be estimated using the signal volume algorithm described above. In the two selected areas shown in Fig. 5b, similar Symbiodiniaceae densities were observed (1.81×10^5^ and 2.07×10^5^ for left and right panels, respectively). The percentage of the hypoxia-stressed Symbiodiniaceae cells differed significantly, from 0.5% (1 out of 196 cells) for the left panel to 44.3% (121 out of 273 cells) for the right panel.

**Fig. 5:**
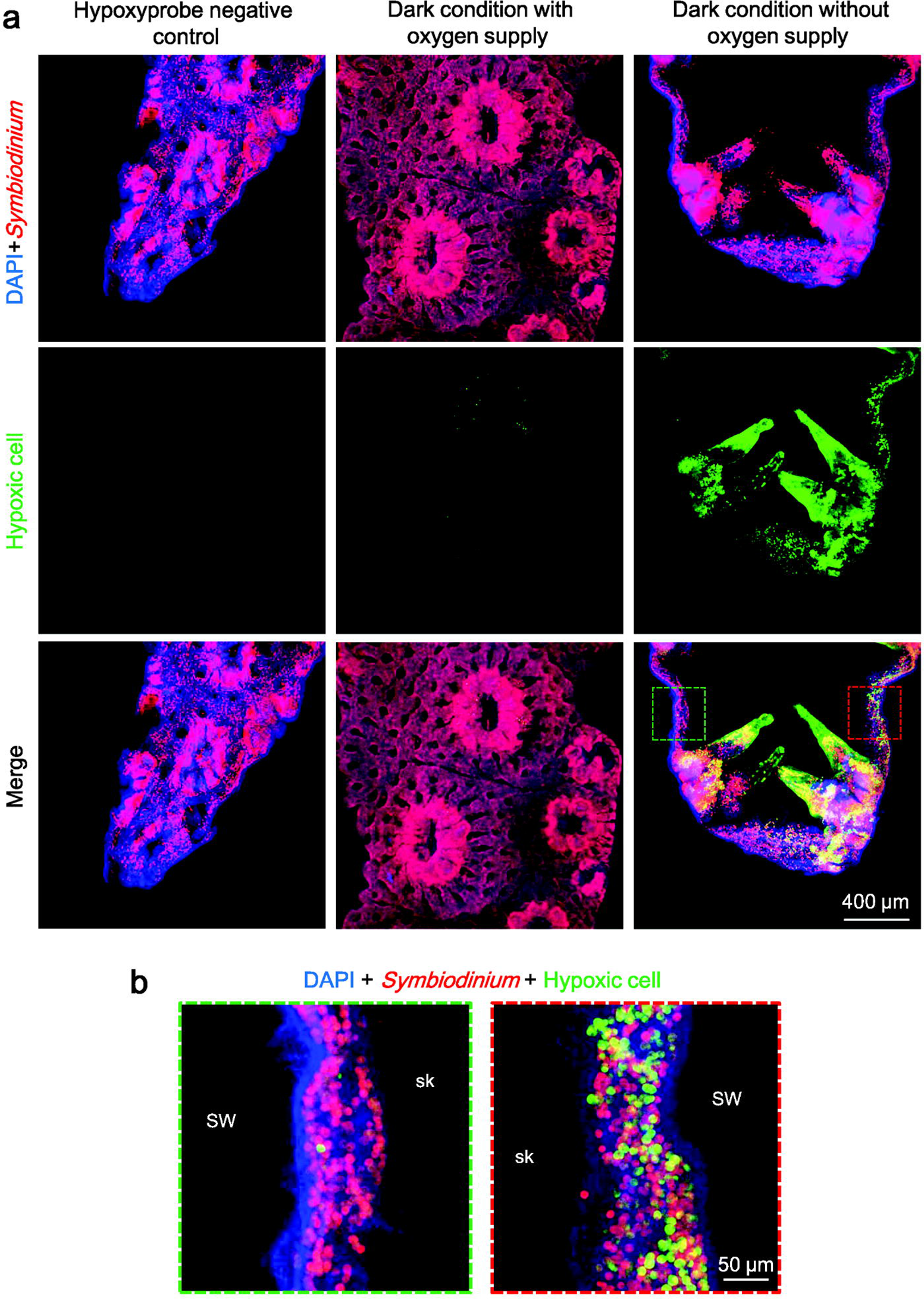
Detection of cellular hypoxia in coral holobionts by using pimonidazole-based hypoxyprobe and TC-LSFM. **a** Immunofluorescence labeling of intracellular pimonidazole (green) in the sample of hypoxyprobe negative control, normoxic control (dark condition with oxygen supply), and the hypoxia treatment (dark condition without oxygen supply). Red depicts the autofluorescence of chlorophylls in Symbiodiniaceae. Blue denotes the DNA-DAPI stain. All of the images were obtained under identical conditions and the same dynamic range and presented as the maximum intensity projection. **b** Increased magnification view of the area marked by the corresponding colored rectangle in the last panel of (**a**), showing the localized distribution of hypoxic Symbiodiniaceae cells in the surface body wall gastrodermis. SW, seawater; sk, decalcified skeleton.

## Discussion

Here, we report the development of a novel technique by combining tissue-clearing technique and light sheet fluorescence microscopy. We also demonstrate that this new technique is powerful for collecting quantitative information on the 3D spatial distribution of Symbiodiniaceae and to elucidate the physiological conditions deep inside the corals with single-cell resolution. The heterogeneous Symbiodiniaceae distribution documented here reveals the superiority of the technique to conventional Symbiodiniaceae density measurement methods. Furthermore, the abundance of Symbiodiniaceae cells in total or in any part of the coral sample, per coral surface area, or per tissue volume could be easily calculated by obtaining the average fluorescence volume per Symbiodiniaceae cell and the total fluorescence volume. Even though fluorescence intensity per cell might change due to physiological changes, our calculation would be unaffected because calculation was based on the volume of fluorescent cells rather than the intensity of fluorescence.

The success in all our initial attempts to couple TC-LSFM with several currently available probes indicates that the proposed technique shows potential for applications that reveal the physiological conditions, including cell proliferation, apoptosis, mitosis, and hypoxia stress, from deep inside coral tissue. Among these conditions, a Symbiodiniaceae mitotic index is an important parameter in determining the symbiosis status of coral holobionts [12,30]. Conventionally, this index is estimated by counting doublet Symbiodiniaceae cells [31], which are actually cytokinetic cells that progress from mitotic cells. In the present study, we demonstrated that immunostaining coupled with TC-LSFM could be applied to accurately estimate the mitotic index. With time-serial sampling, the proposed technique would be instrumental to investigate coral development processes, including metamorphosis of coral larvae and establishment of symbiosis with Symbiodiniaceae, which are critical components of coral population recruitment and growth [32,33].

Hypoxia has been suggested to be an underestimated factor responsible for the decline of some coral reefs [34]. Although photosynthesis by Symbiodiniaceae creates an oxic or superoxic environment during daytime, ambient water can become hypoxic at night, causing stress in corals [28,35,36]. Episodic hypoxia events have been recorded during coral spawning events [37] or phytoplankton blooms [38], eventually causing the extensive mortality of corals and other reef animals in wide areas. Although dissolved oxygen in the gastric cavity of a coral can be measured under laboratory conditions [39], cellular oxygen tension and differential oxygen consumptions by Symbiodiniaceae and by coral hosts have yet to be determined. The gradient of the intracellular hypoxyprobe signal documented here illustrated the diffusion deficiency of oxygen from epidermis to aboral tissue and the respiration of both coral and Symbiodiniaceae cells. With TC-LSFM imaging combined with other techniques, the correlation of cellular hypoxia in coral holobionts with the different genues of Symbiodiniaceae, metabolic condition, apoptosis, and other symptoms of physiological stress could be determined.

TC-LSFM imaging is promising for more studies than shown above. One example is to investigate the distribution and “shuffling” of different genus of Symbiodiniaceae in a coral experiencing thermal or other environmental stress. This technique can be implemented through fluorescence in situ hybridization (FISH) of the sample with a genotype-specific DNA probe, which has been previously coupled with flow cytometry to quantify symbiont assemblages [40]. The spatial differentiation and temporal dynamics of thermal-sensitive and thermal-resistant genotypes can be monitored in situ and in toto in the course of bleaching and recovery, providing data badly needed to test the debated coral adaptive bleaching hypothesis [41,42].

Although some technical challenge remain to be addressed, TC-LSFM coupled with RNA-FISH be used to profile the cell-specific expression of interested genes in a whole coral tissue, which is impossible through qPCR [43], transcriptome profiling [44], or sectioning-coupled RNA-FISH [45].

In addition to the enhancement of data depth and accuracy, TC-LSFM can enable the measurement of multiple parameters on one sample. For example, information on Symbiodiniaceae density, cell proliferation, and apoptosis can be obtained from one sample as long as specific fluorescence probes are available and can be detected through proper filter arrays. With this multiplex approach, correlation between different biological parameters can be achieved with enhanced consistency and reproducibility. Furthermore, this technique reduces the demand for coral samples in research, thereby making it conservation friendly.

TC-LSFM imaging opens up a new horizon in coral biology, and its power to light up the internal world of corals is potentially revolutionary. Coupled with advances in the development of molecular probes and in genomic data [46,47,48,49], the technique can be instrumental for gaining unprecedented depths of understanding on the mechanisms of coral bleaching and resilience under global climate change, and for acquiring information not only of interest to researchers but also fundamental for informed management and conservation of the endangered coral reef ecosystem. Finally, TC-LSFM approach could be adoptable to coral pathology or many other symbiosis cases.

## Acknowledgment

We thank L. Li and W. Yang for their help in underwater sampling and coral aquarium maintenance, respectively. This work was supported by the Natural Science Foundation of China (Grant NSFC 31661143029) and the National Key Research and Development Program of China (Grant 2016YFA0601202 and 2017YFC1404302 for SL); the Research Grants Council Collaborative Research Fund of the Hong Kong Special Administrative Region, China (Project No. 8730037 for SHC); the China Postdoctoral Science Foundation (2017M620271); and the Outstanding Postdoctoral Scholarship, State Key Laboratory of Marine Environmental Science at Xiamen University (for CL).

---

**Supplementary Figure 1 Serial optical sections of the coral samples. a** Optical sections in different *z* axis depths of the 3D imaging of *S. hystrix* branch tip in 100 μm intervals. **b** Optical sections in different *z* axis depths of the 3D imaging of *Seriatopora* sp. polyp in 100 μm intervals. In all of the images, blue indicates the fluorescence of DNA-DAPI stain, whereas red denotes the autofluorescence of chlorophylls in *Symbiodinium* cells.

**Supplementary Movie 1:**

Serial optical sections of *S. hystrix* branch tip in 15 μm intervals.

**Supplementary Movie 2:**

Fly through view of the 3D-reconstructed *S. hystrix* branch tip.

**Supplementary Movie 3:**

Fly through view of the 3D-reconstructed *Seriatopora* sp. polyp.

**Supplementary Movie 4:**

Fly through view of the newly re-attached *P. damicornis* micropropagate.

**Supplementary Movie 5:**

3D reconstruction of *Acropora* sp. spermary.

