## Supplementary figures and images for "Illuminating the dark depths inside coral"

### Supplemental figure 1

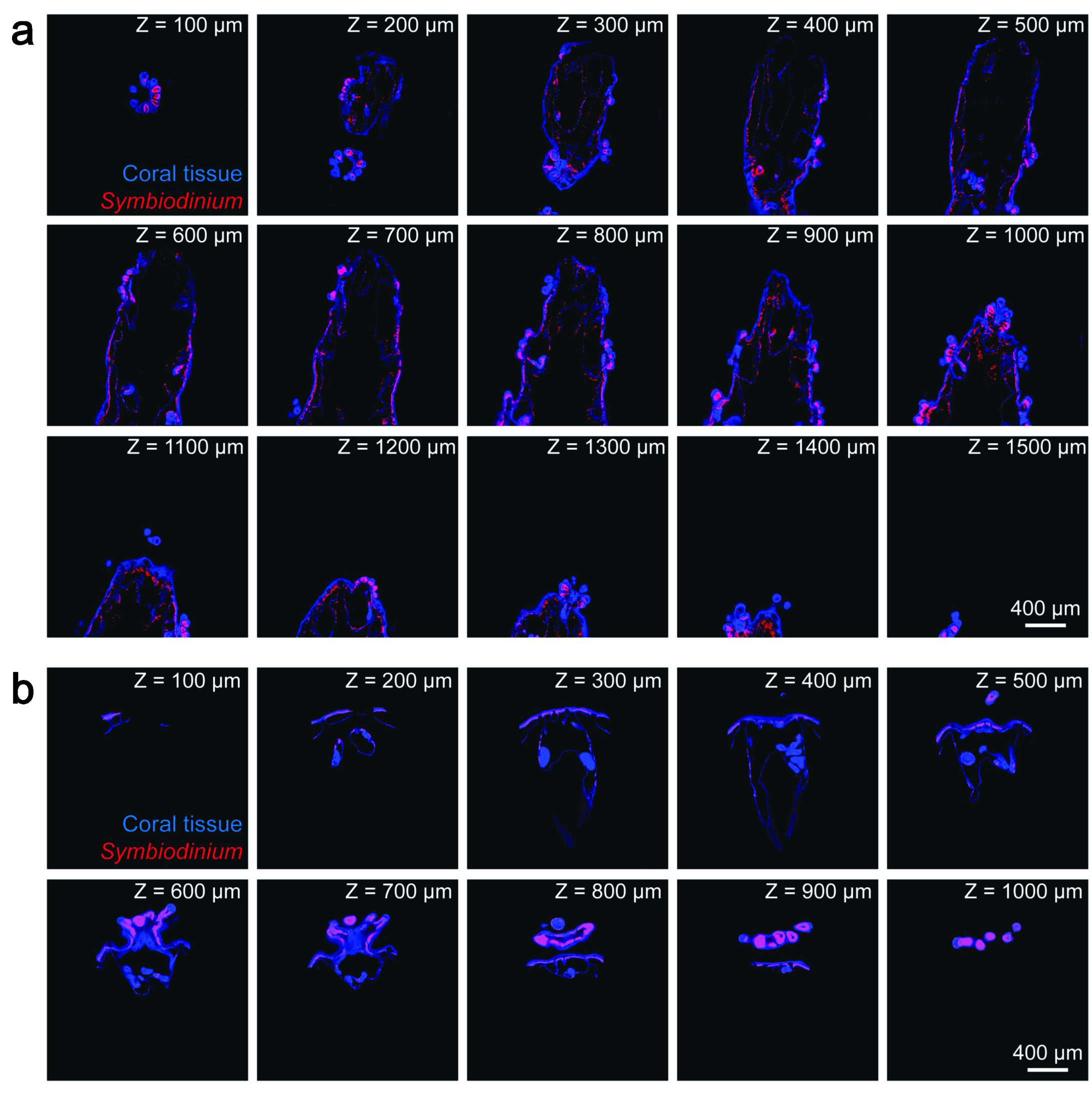
